# Less is More: Oligomer extraction and hydrothermal annealing increase PDMS bonding forces for new microfluidics assembly and for biological studies

**DOI:** 10.1101/150953

**Authors:** L. J. Millet, A. Jain, M. U. Gillette

## Abstract

Key determinants in the emergence of complex cellular morphologies and functions are cues in the micro-environment. Primary among these is the presence of neighboring cells as networks form. Therefore, for high-resolution analysis, it is crucial to develop micro-environments that permit exquisite control of network formation. This is especially true in cell science, tissue engineering, and clinical biology. We introduce a new approach for assembling polydimethylsiloxane (PDMS)-based microfluidic environments that enhances cell network formation and analyses. We report that the combined processes of PDMS solvent-extraction (E-PDMS) and hydrothermal annealing create unique conditions that produce high-strength bonds between E-PDMS and glass – properties not associated with conventional PDMS. Extraction followed by hydrothermal annealing removes unbound oligomers, promotes polymer cross-linking, facilitates covalent bond formation with glass, and retains the highest biocompatibility. Our extraction protocol accelerates oligomer removal from 5 to 2 days. Resulting microfluidic platforms are uniquely suited for cell-network studies owing to high bond strengths, effectively corralling cellular extensions and eliminating harmful oligomers. We demonstrate simple, simultaneous actuation of multiple microfluidic domains for invoking ATP- and glutamate-induced Ca^2+^ signaling in glial-cell networks. These low-cost, simple E-PMDS modifications and flow manipulations further enable microfluidic technologies for cell-signaling and network studies as well as novel applications.

## Introduction

We present a new method for adhering highly biocompatible E-PDMS microfluidic channels to glass substrates that achieves high bond strength without compromising E-PDMS biocompatibility. Microfluidic devices are a mainstay for on-chip miniaturization of chemical and biological systems owing to the highly controlled spatiotemporal manipulations of minute sample volumes. Microfluidics are particularly beneficial for resolving mechanisms of growth, differentiation, and signalling of biological systems that are complex and dynamic. Such microtechnology confers the ability to perform complex environmental manipulations that mimic natural systems on a fundamental level to improve discovery and analysis of biological systems.^1,2^ From manipulations of subcellular domains to miniaturized organs-on-chip, microfluidics facilitate systematic interrogations of cells and networks through the establishment of functional microenvironments.^1,3–8^

The selection and utilization of materials and chemicals are key factors for defining the cellular microenvironment. Over the past century, the means of culturing cells of the nervous system have improved in parallel with the development of a range of platforms and media formulations for chemically-defined investigations into mechanisms of nervous system development and function.^2^ Recent improvements in microfabrication, materials engineering, and chemical processes have provided organs-on-a-chip that refine the ability to achieve biological discoveries and advance medical diagnostics.^3,9–14^

Because of the many advantages inherent in PDMS (transparency, affordability, gas permeability, replication, etc.),^15,16^ it is widely accepted as an ideal material for rapidly fabricating microfluidic devices and environmental culture systems. Thus, it is the most widely used polymer for microfluidic prototyping in research labs and is generally considered to be biocompatible. However, the biological implications of PDMS are process-dependent with conditional biocompatibility that scales with device dimensions and possesses fluidic constraints.^17–21^

Our previous work demonstrates that PDMS can be rendered highly biocompatible, over native unprocessed PDMS or autoclaved PDMS, by extracting the unpolymerized oligomers and metal catalysts from the cured elastomer to produce E-PDMS.^19^ Not only does the solvent extraction process improve the material biocompatibility, but it also alters material properties of PDMS. E-PDMS exhibits reduced adhesion of conformal contact to planar surfaces (e.g., glass coverslips, microscope slides, and Petri dishes)^22^ and increased absorption of small molecules.^23^ While reduced conformal adhesion of E-PDMS confers benefits for substrate patterning and minimizes the transfer of hydrophobic oligomers to the substrate,^22^ E-PDMS provides unique fabrication challenges for implementing microfluidic platforms for cell signalling studies of adherent cells. What is lacking is a thorough understanding of the adhesive properties of E-PDMS and protocols that take advantage of these altered properties.

Typical approaches used to covalently bond PDMS to glass substrates include high-energy processes (plasma exposure, corona discharge, UV-ozone exposure)^24–27^ that have the potential to break bonds and promote the generation of more oligomers. The biological implications of oligomer regeneration in E-PDMS are not well understood. The presence of oligomers in conventional PDMS microfluidic devices has been shown to influence gene expression; even within ethanol-washed PDMS, oligomers still accumulate within cells cultured in the microfluidic channel.^17,21^ Improving highly biocompatible E-PDMS adhesion through non-destructive, bio-compliant processes provides favourable conditions for cell signalling studies in microfluidics and minimizes material-mediated confounds to biological investigations.

Following our previous studies on improved biocompatibility of E-PDMS and substrate patterning with E-PDMS, we observed that cellular processes probe under the E-PDMS to navigate out of the channel between the E-PDMS and glass, indicating a weak and imperfect seal. We also observed distinct increases in bond strengths for E-PDMS microfluidics in conformal contact after long-term cultures (> 10 days), with some E-PDMS microfluidic cultures exhibiting such high affinity for the glass that they could not be removed. Thus, E-PDMS channels in culture either delaminated from the coverslip or they became increasingly adherent the longer they were maintained in the cell culture environment (a humidified atmosphere at 37 °C). We also discovered that by performing solvent-extraction prior to autoclaving, E-PDMS becomes covalently bound to glass substrates. This process also retains the high biocompatibility of PDMS while sterilizing the materials.

Glial cells are the most abundant type of cell in the brain and play important roles in modulating cell communications in the brain. Astrocytic glia sense and respond to local signals, in part through extended branches that contact neighbouring glia, synapses, neurites, and endothelial cells. Conventional cultures in plates, dishes, or wells do not offer the ability to perform selective spatio-temporal interrogation without exposing the population of interest to the stimulant through diffusion. Through microfluidic platforms, signalling between distinct cellular populations can be studied without exposing the entire population to the stimulant. Furthermore, multiple spatial stimulations can be deployed for more complex signalling studies, an experimental advantage not achievable without microfluidics.

Here, we first characterize the bond strengths of E-PDMS as compared to native PDMS over varying regimes of time, temperature, hydrothermal annealing, and glass treatment. We find that, with the exception of plasma treatment that has the undesirable effect of oligomer generation, autoclaving E-PDMS onto acid-cleaned glass produces the strongest bonds. We then exploit these high bond strength E-PDMS microfluidics to compartmentalize primary glial cells and interrogate glial activity by monitoring Ca^2+^ oscillations in response to focal pulses of transmitters applied through laminar flow. Ca^2+^ oscillations are observed in individual cells and waves of Ca^2+^ transients are observed in glial networks. Our work demonstrates the advantage of implementing E-PDMS microfluidic platforms for cell signalling applications.

## Results and discussion

### Materials Characterization

We hypothesized that longer exposure to elevated temperatures and/or humidity facilitates the formation of adhesion bonds between E-PDMS and a clean glass substrate. To test this hypothesis, we fabricated PDMS plugs of uniform volume and geometry and subjected them to various annealing processes on clean glass slides following solvent extraction (Figure 1). We then measured the force required to remove each plug from the glass surface to determine bond strengths. The processes of solvent extraction, or autoclaving, alone improves PDMS biocompatibility and renders the microfluidic environment sterile,^19^ but the combined effects of these processes on adhesion have not been characterized.

**Figure 1.**
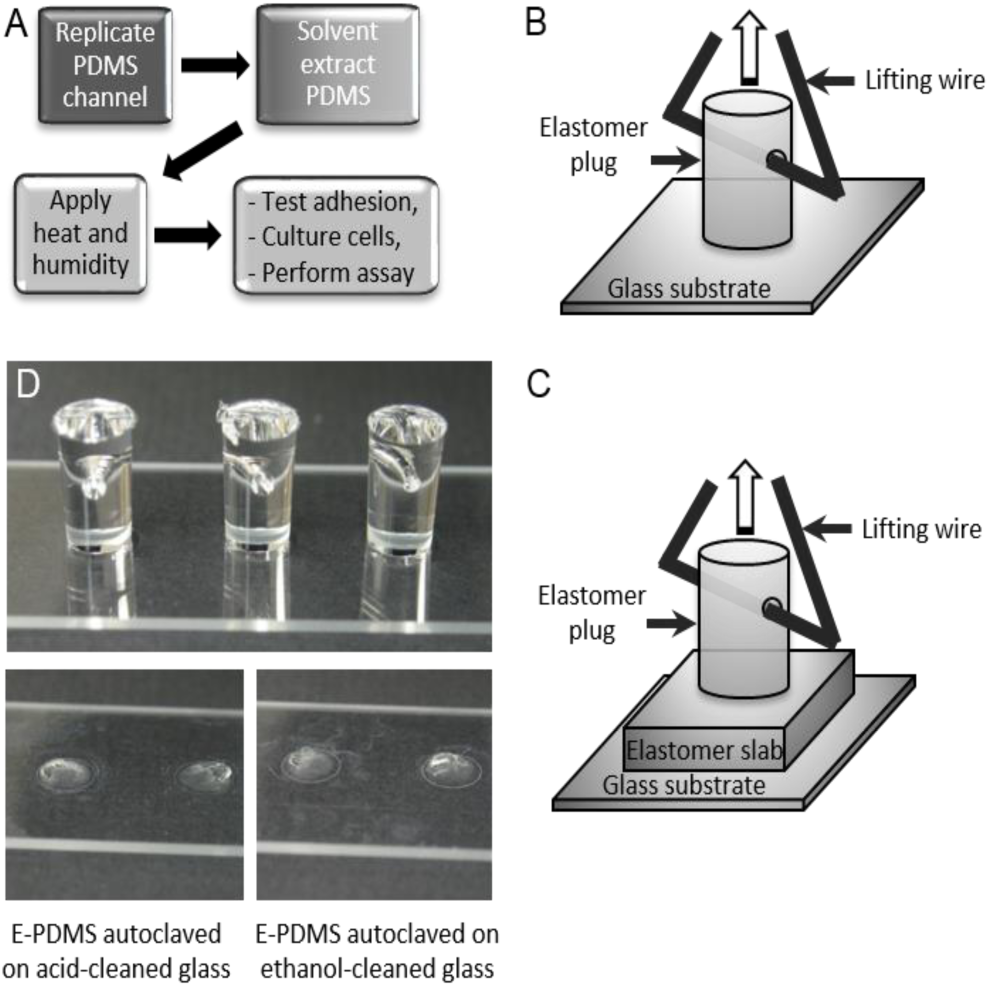
Overview of E-PDMS annealing process and adhesion measurements. (A) General overview of annealing process, from replicate moulding of PDMS through cell culture. For microfluidics, masters are used to generate the PDMS microchannels. For plugs to measure adhesion forces, a 96-well plate serves as the replicate mould. Solvent extraction was performed with n-pentane, xylene, ethanol (200 proof), and water. (B, C) Schematic of PDMS elastomeric plugs and substrate configurations. Holes punched through the plugs allow for wire supports to attach to the force scales. PDMS on glass (B) was used for annealing measures. PDMS on PDMS (C) was used to measure material deformations through profilometry. (D) Images of 3 plugs on a microscope slide for force measurements (upper image). E-PDMS plugs form strong adhesion bonds after solvent extraction and autoclaving. It is typical for E-PDMS annealed after extraction and autoclaving to tear, leaving elastomer fragments when being pulled off the glass, irrespective of glass cleaning method (lower images).

Figure 2 shows a time-dependent increase in normal bond strengths of E-PDMS plugs on glass slides exposed to a range of conditions. For cell culture conditions (Figure 2A), there is a significant increase in the adhesion bonds (from 65.1 to 286.6 kPa), (p < 0.02, two-way ANOVA) of E-PDMS on acid-cleaned glass when exposed for 2 weeks to a humidified atmosphere at physiological temperatures (37 °C). Our glass cleaning process for primary cell culture uses a sulfuric acid bath followed by a DI water wash. A common alternative cleaning protocol is sterilization with ≥ 70% ethanol in DI water. We included both glass preparations in our E-PDMS and PDMS adhesion force comparisons. Our results show that under cell culture conditions, acid-cleaned glass enables a stronger adhesion of E-PDMS to glass than does the ethanol-cleaned glass.

**Figure 2.**
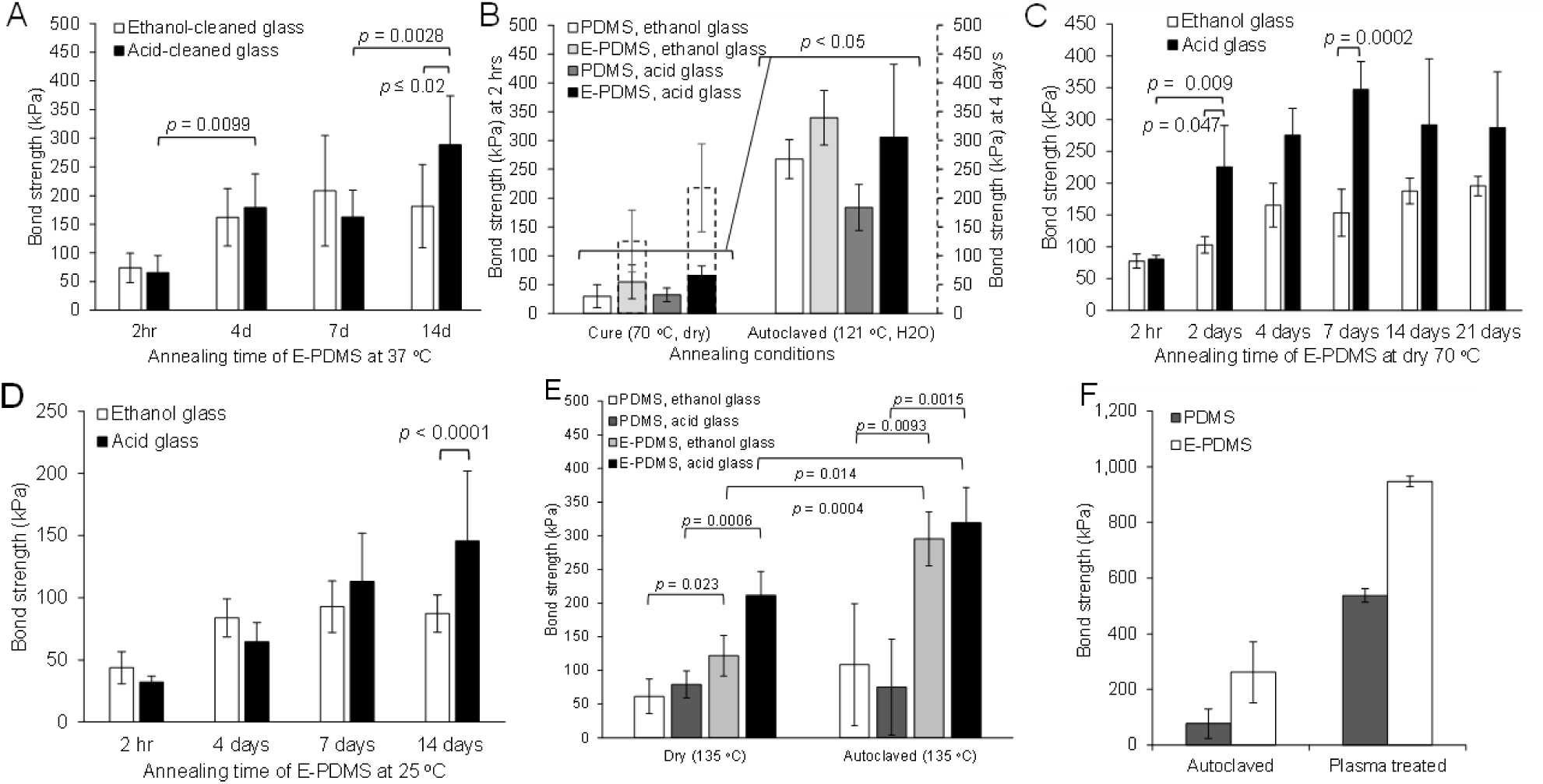
Measurements of removal forces for annealed E-PDMS and PDMS plugs. (A) Forces required to remove annealed (humidified 37 °C) E-PDMS plugs off microscope slides were measured. Similar results are achieved for both ethanol and acid-cleaned glass at 37 °C. (B) Significant temporal effects of two annealing conditions were tested for both PDMS and E-PDMS plugs on ethanol-cleaned glass (white and light grey bars) and acid-cleaned glass (dark grey and black bars). E-PDMS annealed with dry heat (4 days) has a significant effect on changes of removal forces. Autoclave annealing produces superior bonding forces. (C) Temporal-dependence of annealing E-PDMS with 70 °C dry heat. (D) E-PDMS plugs retained at ambient temperature and humidity show increased removal forces over a two-week period. (E) Measurements of high temperature-matched annealing (135 °C) under dry or humidified conditions show that heat, humidity, and E-PDMS produce the highest bonding forces. (F) Comparison of extraction-autoclave annealing to plasma-heat annealing. E-PDMS out performs PDMS in both cases (n=4, One-way ANOVA, p<0.05). (A-E) n=8, Two-way ANOVA.

To resolve the contributions of conventional fabrication and sterilization conditions to that of increased E-PDMS bonding strengths, we compared E-PDMS annealed to glass with dry heat (PDMS curing oven, 70 °C) vs. humidified heat (autoclave sterilization, 121 °C). Figure 2B shows that autoclaving increases adhesion of E-PDMS and PDMS over dry heat exposure for both material types (p < 0.05, two-way ANOVA). E-PDMS showed superior adhesion over PDMS, regardless of treatment. After 4 days exposure of E-PDMS and PDMS to dry heat, PDMS adhesion results were not statistically different from 2 h, whereas E-PDMS on acid-cleaned glass over 4 days showed a statistically significant increase (from 65.1 kPa to 218.2 kPa, p < 0.001, two-way ANOVA); the increase on ethanol-cleaned glass at 4 days was not significant.

We performed an extended time-course study on the temporal-dependence of E-PDMS annealing under dry heat (Figure 2C). The data show that after 7 days at 70 °C E-PDMS normal bond strengths (153.1 to 348.5 kPa) are significantly greater on acid-cleaned glass than the alternative (p = 0.0002), and normal bond strengths become similar to those achieved through the 2-h autoclaving process (65.1-348.5 kPa) (Figure 2B). These data suggest that increased heat promotes the annealing of E-PDMS to glass. E-PDMS plugs kept at room temperature (25 °C) for 14 days did not show this large of an increase in normal bond strengths but at 14 days had significantly stronger (87.9 kPa verses 146.6, p < 0.0001, two-way ANOVA) bonding than PDMS (Figure 2D). To further understand the influence of temperature and humidity, we exposed both E-PDMS and PDMS plugs to temperature-matched (135 °C) treatments in a dry oven and in an autoclave with saturated humidity (Figure 2E). Autoclaving produced significantly higher normal bond strengths than dry heat for E-PDMS on acid-cleaned glass (p = 0.014) and ethanol-cleaned glass (p = 0.0004). Again, autoclaving provides superior results for E-PDMS, suggesting that both humidity and elevated temperatures facilitate the annealing process of E-PDMS to glass. It should also be noted that it is possible for PDMS and E PDMS structures to have altered dimensional stability and tear strength after autoclave sterilization^.28^

For perspective, we compared our E-PDMS annealing process to the typical oxygen plasma-bonding process. E-PDMS material can be activated through high energy processes; however, these high energy processes break PDMS bonds and introduce more oligomers into the culture system.^24,25,27^ Our results show that E-PDMS adhesion outperforms PDMS after autoclaving or plasma treatment, and plasma treatment shows superior normal bond strengths compared to autoclaving (Figure 2F). It remains uncertain how much oligomers are generated in E-PDMS through high energy processes, such as plasma treatment, or how freely any newly formed E-PDMS oligomers can translocate from the material to cells cultured in microfluidic systems. From these data, it is clear that solvent-extraction of unbound oligomers and catalyst from PDMS increase bonding forces through a thermal hydration process. It is known that siloxane and silanol groups equilibrate with a humidity-dependence; silanols are very unstable and the reaction is highly skewed in favour of forming thermally stable siloxane bonds through a condensation reaction.^29–32^ Nevertheless, hydrogen bonding, van der Waals forces, and nano-scale interlocking interactions of the glass and PDMS cannot be underestimated and could also contribute to an increased adhesion in addition to siloxane bond formation. Even the inclusion of silica filler particles in PDMS can influence the material properties, thus having further implications for PDMS extraction.^33^ Given these adhesion study results, we employed solvent extraction followed by autoclave annealing for microfluidic cell cultures with fluid manipulation for cell signalling studies.

With the ability to covalently bond E-PDMS to glass through an autoclaving process, we tested the possibility of performing polymer-to-polymer bonding through extraction and autoclave annealing. We placed E-PDMS and PDMS materials each in direct contact on both substrates of E-PDMS and PDMS, then subjected them to the same autoclaving process. Following the annealing process, we observed a consistent trend of noticeable location-dependent deformation of the plug “footprint” when plugs were placed on the opposite substrates and autoclaved. Profilometric measurements show that PDMS substrates indent (~7 μm deep depression) where E-PDMS plugs are placed in direct contact during autoclaving (Figure 3). Conversely, the footprints on E-PDMS substrates swell (~6 μm vertical change) only where PDMS plugs are in contact with E-PDMS substrates (Figure 3B). The footprint deformation induced by autoclaving E-PDMS in contact with PDMS is significant (p < 0.0001) and consistent regardless of whether E-PDMS serves as the plug or substrate (Figure 3C). No appreciable changes of footprint deformation are observed when E-PDMS and PDMS plugs are matched to the self-same substrates.

**Figure 3.**
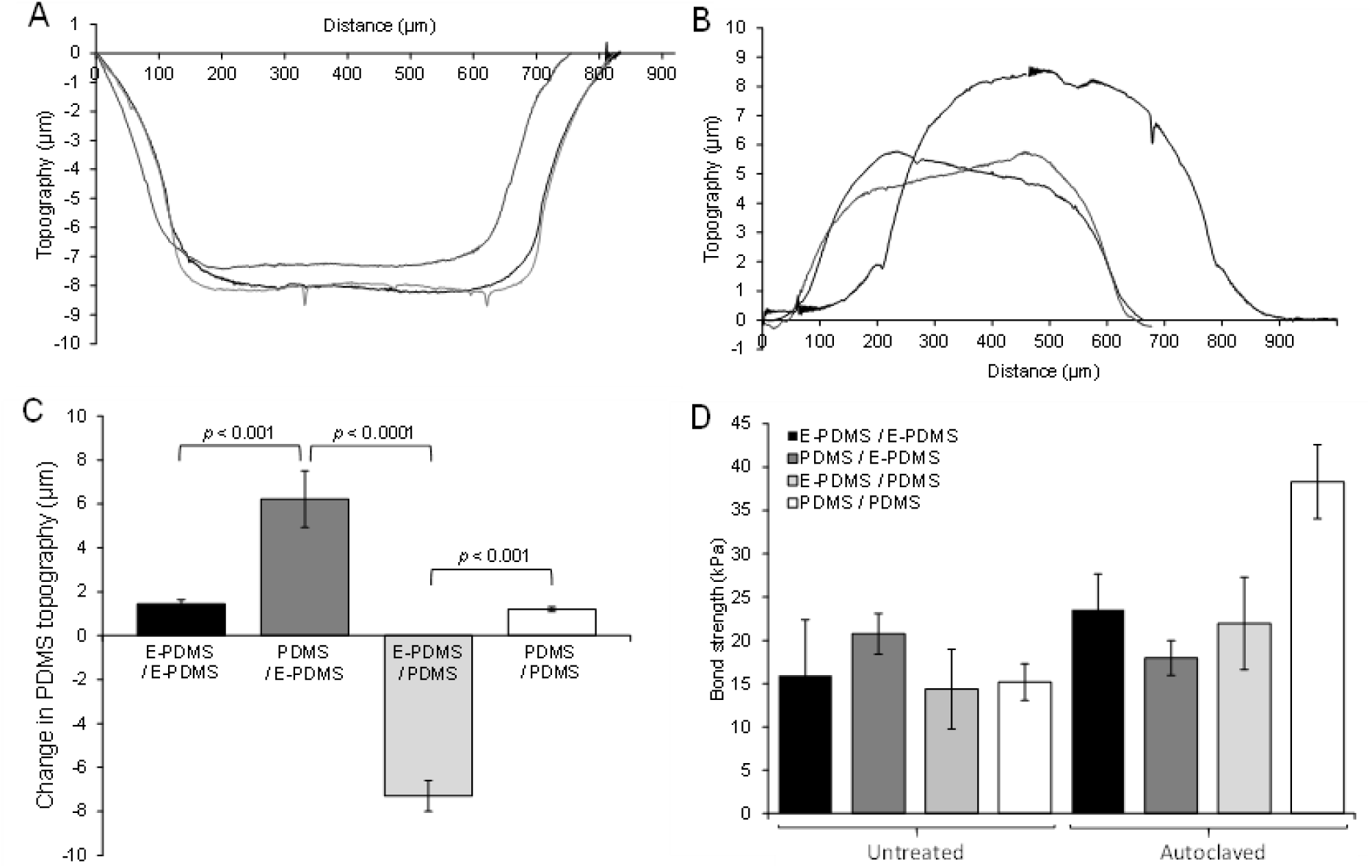
Measurement of deformation and removal forces for E-PDMS and PDMS in direct contact. (A, B) Profilometry data confirms the visual observations and profiles topological changes of E-PDMS and PDMS deformation when extracted and non-extracted samples in direct contact are autoclaved. (C) Results of changes in topography from profilometry data of E-PDMS and PDMS plugs in contact with the E-PDMS and PDMS slabs. PDMS/E-PDMS combinations show much greater changes (p < 0.001, 3 repeats (n=3 each), One-way ANOVA) than both E-PDMS/E-PDMS and PDMS/PDMS; matched material types show no significant changes. (D) Bond strength measurements of autoclave-annealed E-PDMS and PDMS plugs on E-PDMS and PDMS slabs. Weak bond strengths for PDMS on PDMS are higher than any E-PDMS interactions and are attributed to the remaining oligomers and metal catalysts left in the bulk polymer (no significance).

Oligomer translocation is possible in cross-linked PDMS as evidenced by the time-dependent reversion of plasma treated PDMS from a hydrophilic to hydrophobic state.^24,25,34,35^ This information, combined with our results of E-PDMS and PDMS deformation, suggest that oligomer translocation occurs within PDMS and the effect on the material is measurable after translocation from source (PDMS) to sink (E-PDMS). The observable deformation indicates that a pronounced quantity of material remains unpolymerized when prepared under common conditions (10:1 ratio, 70 ^oC^ cure for 2 hrs). It also provides further evidence to support the need to remove unpolymerized oligomers through the pentane-xylene-ethanol-water solvent extraction process.

Bonding force measurements show that PDMS-PDMS pairing achieves the highest bonding strengths (38.4 kPa) over any of the other pairings of the two polymer types (Figure 3D), possibly due to the presence of oligomers and the platinum catalyst that can facilitate further chemical interactions between untreated PDMS materials at elevated temperatures. Without oligomers and catalyst in one of the two paired materials, bonding strengths are on average half (12.7-25.4 kPa) those achieved with PDMS-PDMS interfaces. Solvent extraction removes oligomers and platinum catalyst from the polymerized material,^19^ thus reducing its ability to crosslink with other PDMS surfaces but improving the ability of E-PDMS to bind to glass. These results provide new data and perspective that adds further evidence to the understanding that E-PDMS possesses unique material properties, beyond increased biocompatibility, apart from its native, unextracted material form.

### Cell Signalling

With the ability to produce highly-biocompatible E-PDMS tightly bound to clean glass substrates, we cultured primary astrocytes in E-PDMS microfluidic platforms for cell observation and analyses. These platforms enable spatiotemporal manipulation of fluids to achieve multiple, simultaneous or sequential focal stimulations across compartmentalized cultures. Following solvent extraction, E-PDMS microfluidics were trimmed to produce fluid source wells useful for retaining and perfusing media during cell culture, for actuating fluids during cell signalling studies, and for immunolabelling cells (Figure 4). Fluidic pulses, achieved through a combination of gravity and hydrostatic pressure, were produced in adjacent channels of the microfluidics to demonstrate a range of fluidic controls from seconds to minutes (Figure 5). The method of chemo-temporal manipulation employed enables rapid chemical transients without increased pressures within the microfluidic device.

**Figure 4.**
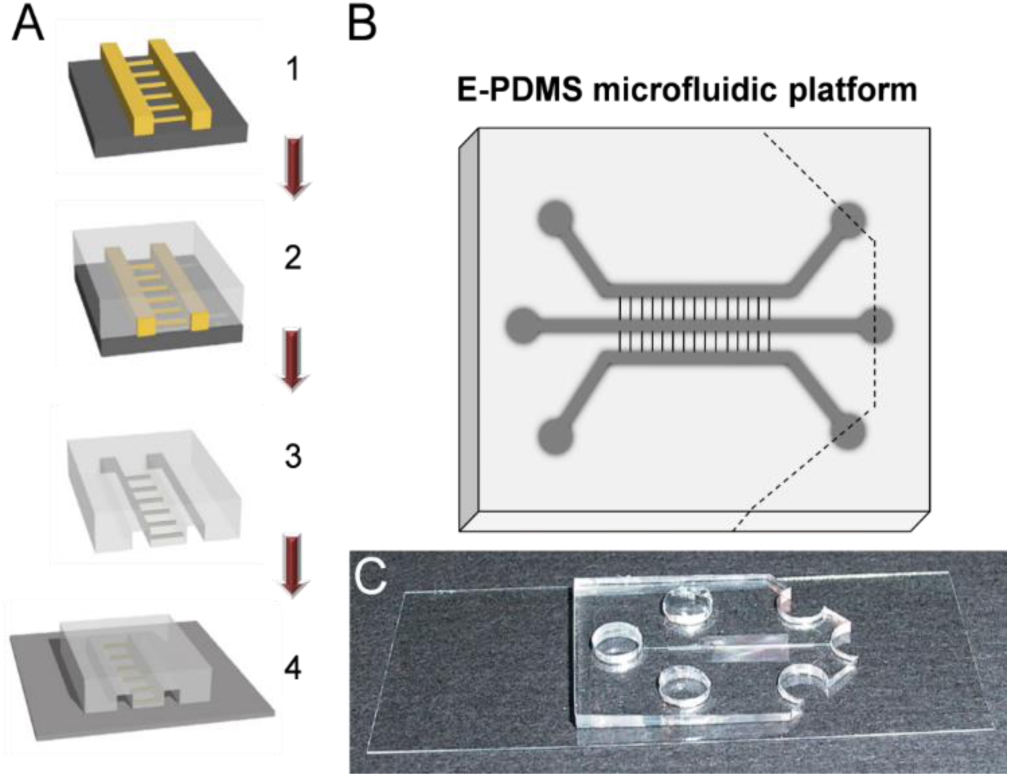
Assembly of microfluidic devices for cell signalling studies. (A) Overview of fabrication and assembly process for microfluidic chambers (200 μm W x 45 μm H) for cell signalling. (1) Microfluidic masters are used for (2) replicate moulding of PDMS-based microfluidic channels with interconnecting tunnels (7x7x45 μm). (3) PDMS replicates are trimmed and holes punched (upper right diagram) prior to solvent extraction to remove un-cross-linked oligomers and metal catalysts. (4) E-PDMS replicate is annealed to the glass substrate through autoclave sterilization, completing the microfluidic device. (B-C) Architecture (B) and photograph (C) of an annealed E-PDMS microfluidic device used for cell signalling studies. Ports are cut with dermal biopsy punches (5-6 mm). The reservoir is made by bisecting three ports on one end of the platform (dashed line).

**Figure 5.**
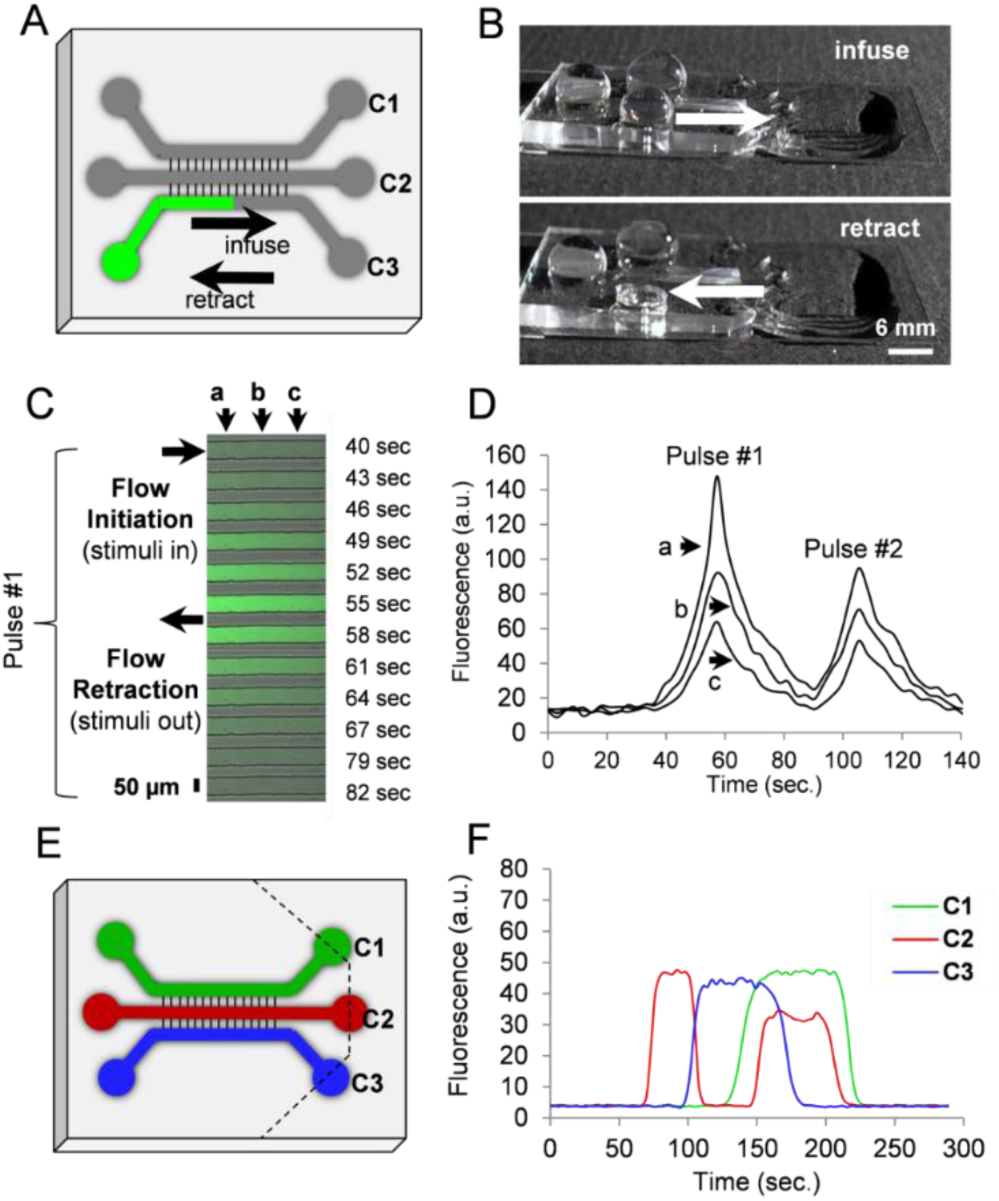
Chemotemporal pulses in microfluidic channels. (A) Schematic and image of PDMS channels on a coverslip for a fluidically-connected microfluidic platform for cell signalling. (B) Using surface tension and gravity flow, chemicals can be pulsed in the channel for stimulating compartmentalized cells. (C) For pulse characterization, dilute FITC-conjugated secondary antibodies (green) were introduced in the channel. Spatial characteristics were assessed at three separate points 125 μm apart (a, b, c). (D) Two serial pulses demonstrate spatio-temporal characteristics for multiple cellular stimulations. In pulse #1, half-maximal stimulation concentration is achieved at peak (a) with a concentration decay of 40% at peak (b), and 57% at peak (c). Positions of points a, b, c as in (C). (E) Parallel pulsatile flow actuated by surface tension and gravity-mediated passive pumping within the same device. The device was cut along the dashed lines to allow for unrestricted outflow. Pulsatile flow was accomplished as shown in (B) with an increased fluid head at the inlet initiating channel infusion, and emptying of the inlet resulting in flow retraction. (F) Flow in the three individual channels (C1, C2, C3) can be modulated independently to produce spatio-temporal pulses as depicted in the profiles of fluorescence. Flow was initiated in C2, then retracted in C2 at the same time as C3 initiation; infusion of C1 and re-infusion of C2 precede retraction at C3, ending with serial retractions from C2 and C1.

This approach to fluidic actuation permits simultaneous pulses in multiple channels, is applicable for any lab seeking to employ microfluidics, and does not require expensive pumping systems. Like neurons, glial cells can be guided into adjacent microchannel compartments. Individual glial cells cultured in E-PDMS microfluidics develop glial extensions that migrate along channel corners to extend through 3 μm interconnects. Most glia show affinity for the glass-to-sidewall channels interface. Larger ramified glial cells do not prefer channel corners (Figure 6) but appear to avoid them. Similar observations of corner affinity, or avoidance, have been noted for other cell types in microfluidics.^19,36^

Glial protrusions possess sensory capabilities for detecting changes in extracellular concentrations of small signalling molecules (e.g., ATP, glutamate). Glial cells exhibit spontaneous or evoked fluctuations of internal Ca^2+^ stores that can induce propagating signals that pass through nervous tissues and cell cultures.^37–39^ Potential roles of Ca^2+^ waves include contributions to vascular regulation, metabolic processes, synaptic modulation and mechanisms of axonal guidance.^40–42^ Ca^2+^ transients in glial cells exhibit characteristic temporal signature forms and periodicities.^43–45^

**Figure 6.**
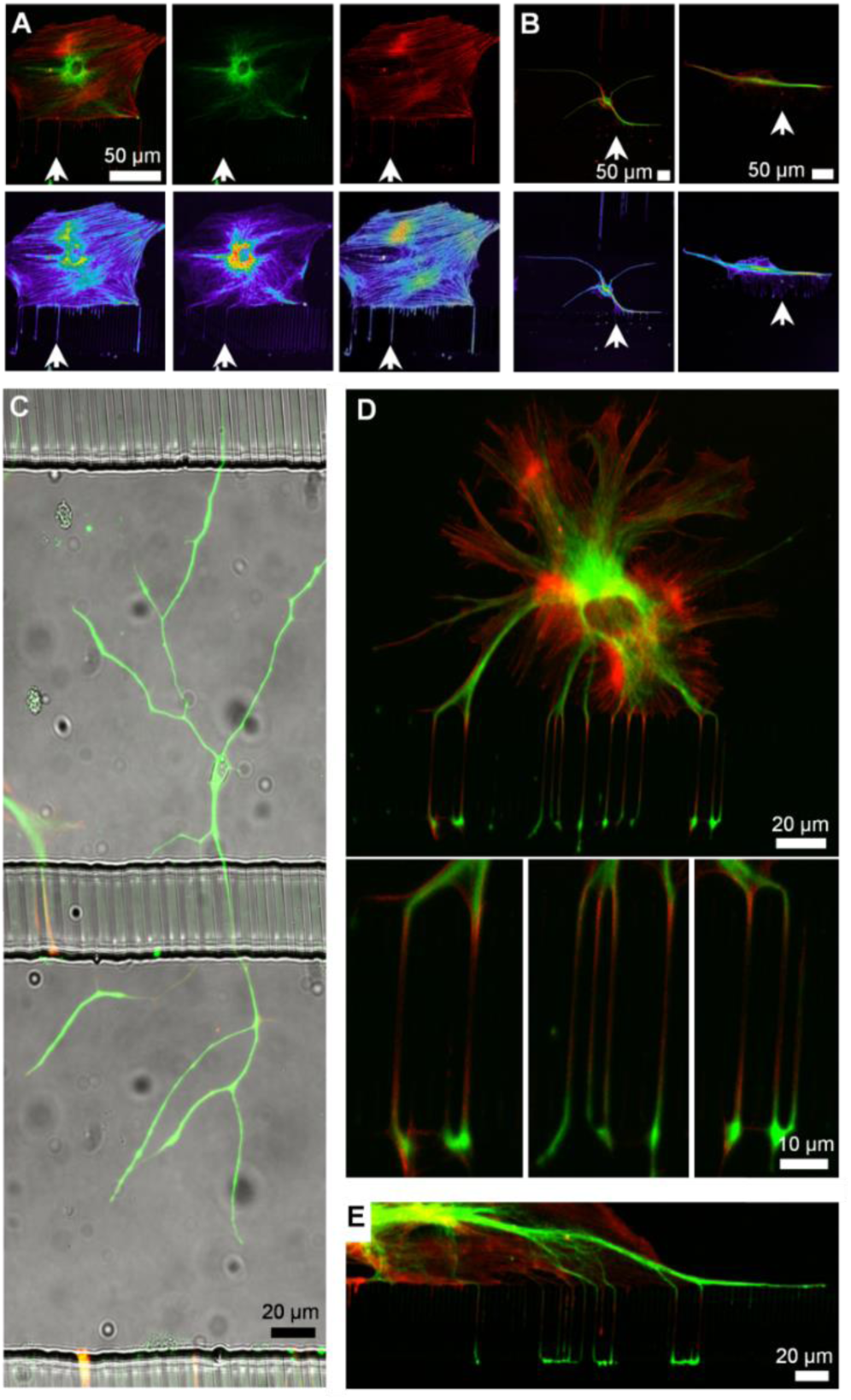
Compartmentalization of glia in microfluidic channels. (A, B) Glial cells cultured in contact with E-PDMS microfluidics develop cellular branches that extend through interconnects. Antibodies for glial fibrillary acidic protein (green) and rhodamine phallotoxins (phalloidin, red) label the cytoskeleton. Fluorescence intensities of cell branches are represented by glow-scale intensity images (below). Fine, filamentous glial branches are rich in filamentous actin and contain GFAP filaments. (B) Merged fluorescence images of two glial cells are shown, each in its own image pane. (C) Large (~400 μm long), ramified glial cells do not show affinity for channel side-walls and interconnects as most glial cells do. (D) A glial cell in the top channel extends branches into the bottom channel allowing subcellular stimulation with signalling molecules applied through adjacent channels. Insets show magnified views of glial branches wrapping around the interconnect pillars, merging as they emerge into the bottom channel. (E) Glial cells often spread along the interconnect wall sending branches over distances exceeding 100 μm, increasing the efficiency of inter-channel signal transmission.

Figure 7 shows a population of glial cells stimulated through a subcellular plug of ATP (10 μM) focally administered to a compartmentalized cell. Within the population, some glial cells show brief, robust cyclic or acyclic response, while other cells continue to oscillate beyond the 4 min observation time. Previous studies have typified Ca^2+^ responses with high throughput analyses; signals of glia in E-PDMS microfluidics conform to categorical morphologies of spikes, bursts, cyclic responses, or sustained Ca^2+^ elevations.^43^ The rate of Ca^2+^ wave propagation (11-12 μm/s) for glia cultured in our E-PDMS microfluidic samples are within the range of wave propagations observed in dispersed cultures and brain slices (6-27 μm/s).^46^ The process of extracting unpolymerized oligomers and binding E-PDMS to the glass through autoclaving both removes harmful oligomers that can accumulate in cells.^19,21^ and provides strong normal bond strengths that allow for cell signalling studies without E- PDMS delamination. With the ability to develop and stimulate compartmentalized glial cultures without biasing the remaining culture to stimulating chemical cues, we cultured glial populations in the central channel between two parallel microfluidic stimulation channels.

**Figure 7.**
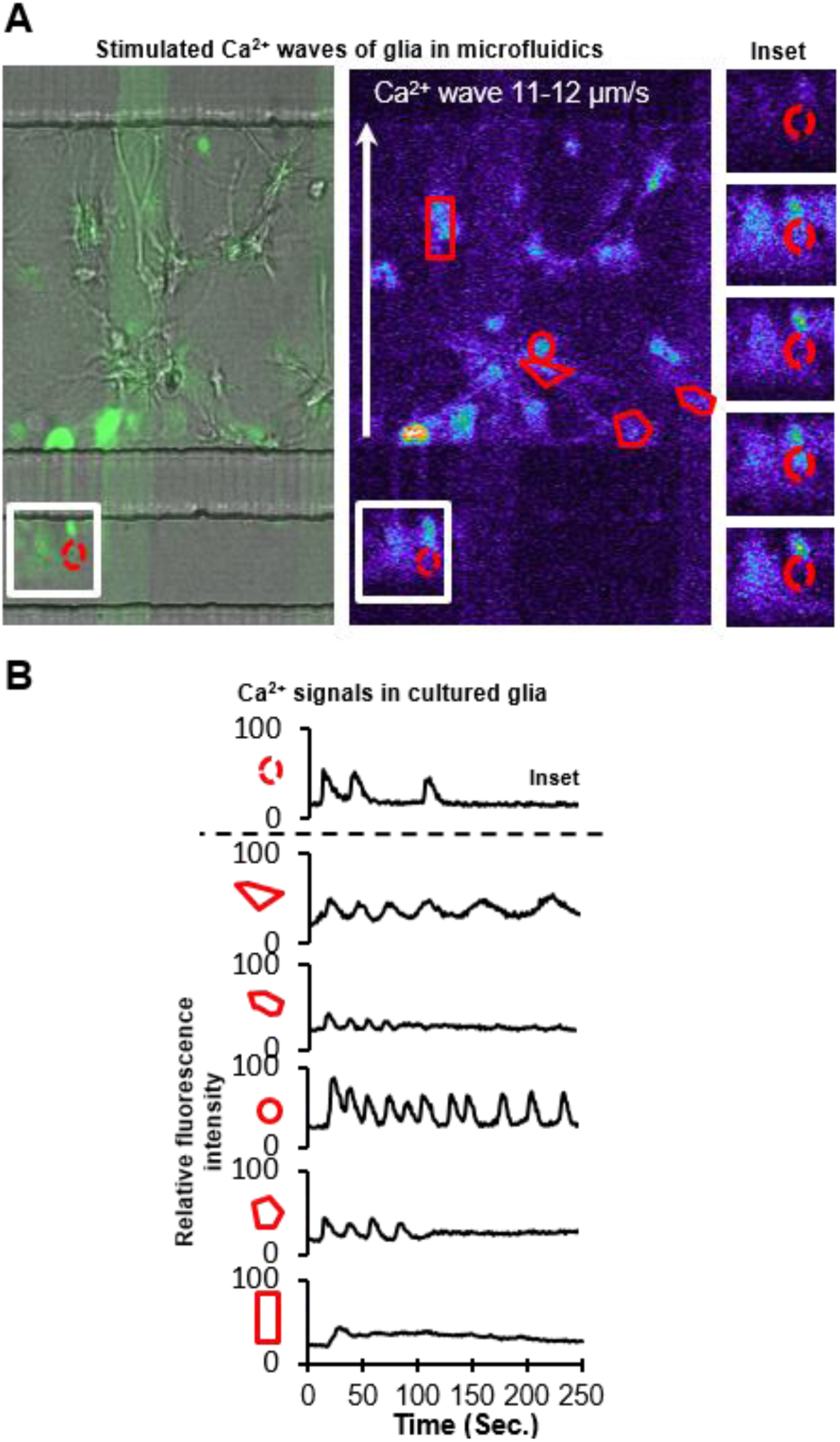
Ca^2+^ signals of glial networks in E-PDMS microfluidics. (A) Primary glial cells proliferate and extend branches into adjacent microchannels through interconnects. All channels were loaded with Ca^2+^ indicator (Fluo-4 AM in PBS). Glial cell branches were stimulated via 10 μM ATP infusion (left to right) through stimulation channel (bottom). A combined brightfield and fluorescence image is shown (left) alongside the respective fluorescence intensity glow-scale image (right); FITC-PLL lines (green) on glass are reference markers. Segmented images of inset (right) demonstrate repeated Ca^2+^ fluctuations. (B) Fluorescence profiles of Ca^2+^ responses from respective ROI demonstrate the range of characteristic Ca^2+^ responses for cultured astrocytes. Glia cultured in microdevices exhibit a typical range of Ca^2+^ fluxes that vary in frequency, amplitude, and duration.

Glial cells develop in the central channel and expand in clusters to extend processes into adjacent channels throughout the length of the microfluidic platform. ATP (20 μM) was administered before glutamate (50 μM) in quick succession (73 s between applications) (Figure 8). Although two inputs were applied, three distinct population responses were observed (Figure 8A-B). First, ATP created a local response on one side of the population. A second stimulation locally activated the opposite portion of the population, followed by a third robust response on the same side as the second response. The third stimulation induced activation of the entire population, reminiscent of a microburst (Figure 8B). While the source of the additional Ca2+ response is not known, the most plausible explanation would be that the ATP plug activated an upstream glial population, glial processes extending into the second stimulation channel may have released a chemical messenger into the channel prior to the introduction of the glutamate application. Through laminar flow, the chemical messenger could be carried down the channel to activate downstream glial populations. Figure 8C schematic summarizes the sequence of cellular “regions of interest” activated and displayed in Figure 8B.

**Figure 8.**
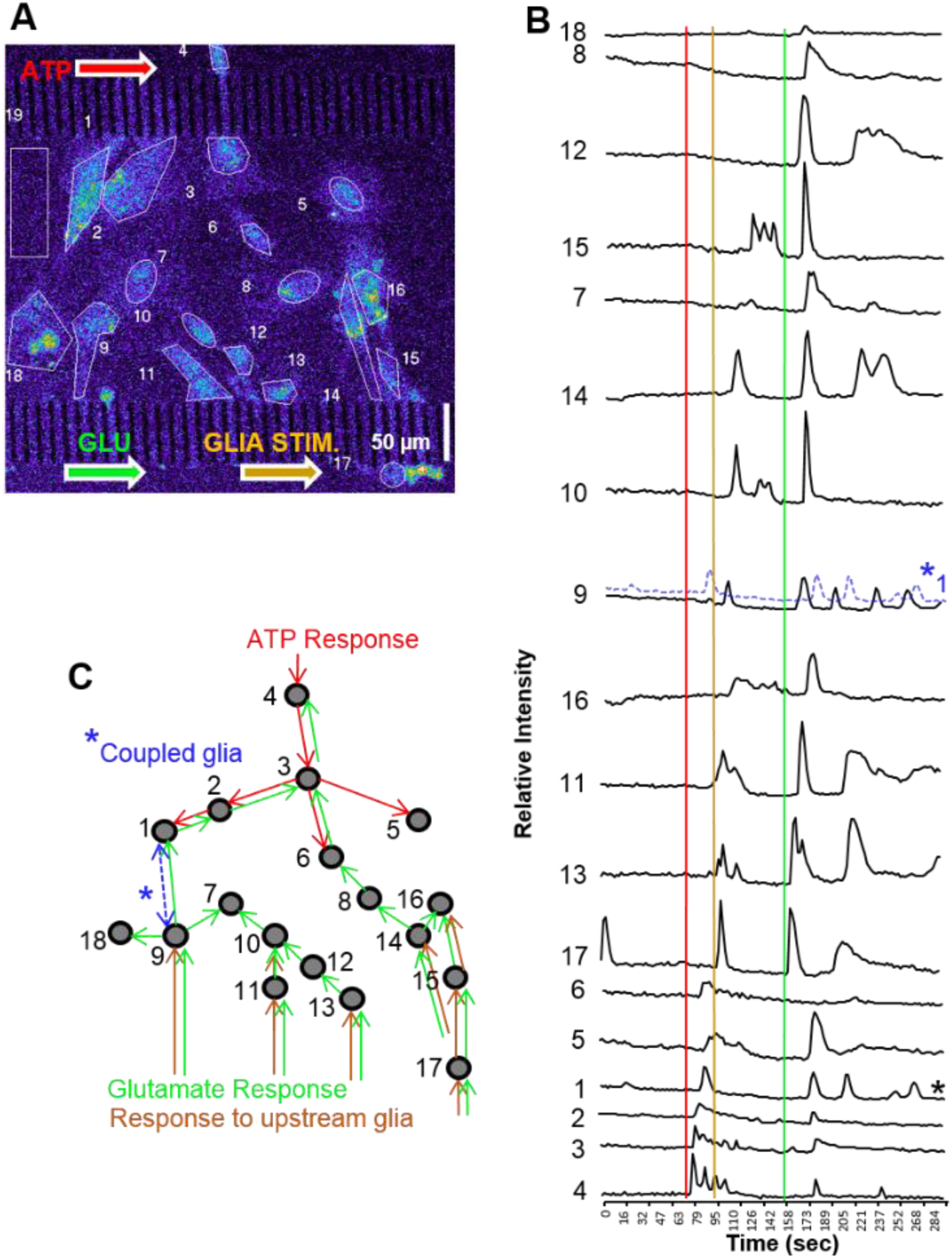
Glial network activation through multipoint stimulation. (A) Glia were cultured in the central channel of a microfluidic platform for stimulation at opposite sides of the culture for observing network activity. First, 20 μM ATP was introduced, then 50 μM glutamate was inserted into the bottom channel. From two chemical stimulations, three network activations occur. (B) Background (ROI#19)-subtracted fluorescence intensity profiles of each defined RIO ordered in chronological activation. Temporally-coupled signals from cellular ROI 1 and 9 suggest the two cells are electrically coupled. (C) Schematic summary of Ca^2+^ signal propagation through the glial population.

Glial cells are known to signal both through gap junctions and vesicular release.^40,47^ Gap junctions permit local signal transfer to directly connected cells whereas gliotransmitter release can influence cells not connected through gap junctions.^48^ We demonstrate the ability to culture glial cells in E-PDMS microfluidic devices for cell signalling investigations. A handful of studies co-culture glial cells with neurons in microfluidics for studying neuro-glial interactions.^49–54^ To our knowledge, no investigations specifically study glial population signalling interactions in microfluidic devices.

Microfluidic environments surpass cell culture dishes in their ability to exert spatio-temporal control of the microenvironment. Glial cells stimulated in a dish can release gliotransmitters that diffuse from sites of stimulation to neighbouring cells. Culture-dish perfusion chambers allow for continuous fluidic exchange that can rapidly wash stimuli away; however, the entire culture is exposed to flow conditions that carry the stimuli across the remaining population or dilute chemical messengers released locally. Microfluidic environments overcome many of the limitations of dish cultures, perfusion chambers, or in vivo, while enabling signalling studies not previously achievable.

## Experimental

### Preparation of PDMS plugs

PDMS plugs were prepared by mixing prepolymer and curing compound (Sylgard 184, Dow Corning) at a 10:1 ratio, respectively. After thorough mixing, all wells of a clean 96- well plate were filled with the PDMS mixture and cured at 70 °C for at least 2 h. Care was taken to prevent filling voids between wells with PDMS. A small amount of 70% ethanol in deionized water (DI, from a Millipore (MilliQ) filtration system) was used to facilitate releasing cured PDMS plugs from individual wells. PDMS plugs were dried of ethanol, then a 1.0 mm dermal biopsy punch was used to bore a diametric hole through the vertical midpoint of the plug. PDMS plugs were separated into two groups for treatment (solvent extraction) and control group (no treatment) prior to assembling plugs onto microscope slides or PDMS slabs.

### Preparation of Microfluidic Channels

Microfluidic channels were replicated using the same master sets from our previous work for substrate patterning; for master fabrication and PDMS microchannel replication from a library of devices, we refer the reader to our initial description.^22^ The cured PDMS microfluidic channel replicates were solvent extracted through a simplified protocol (protocol defined below) followed by a thermal bonding process using an autoclave.

### PDMS Extraction

Prior to assembling PDMS plugs or channels onto slides or coverslips, E-PDMS structures were extracted through a series of organic solvents according to the optimized extraction protocol modified from our original process,^19^ while the control group was left untreated. For microfluidic channels, four to five PDMS microchannel replicates measuring ~464 mm^2^ and 2–4 mm thick were gently stirred and submerged in 150–200 mL of each solvent for the indicated times: HPLC grade pentane (Fisher Scientific) for ~16 h; xylenes isomers with ethylbenzene 98.5+% (xylenes) (Sigma) for 1–2 h; xylenes for 2–4 h; 200 proof ethanol (EtOH) USP for 1–2 h (AAPER); EtOH for 2 h minimum.

In this protocol, pentane is used to swell the PDMS, xylenes are used to remove the pentane, ethanol is used to remove the xylene, water removes ethanol. This gradual solvent exchange returns the PDMS to its former size. In the final step, the PDMS channels were transferred from ethanol to 1 L of sterile DI water and soaked overnight, then dried prior to use. For E-PDMS plugs, the same solvent extraction sequence was employed using 500-600 mL of each solvent. To ensure the PDMS plugs were devoid of any residual solvents, plugs were dried at least 1 week (ambient temperature and pressure) after the extraction process, prior to use.

### Glass Cleaning

Two common methods of glass cleaning were employed for comparison, ethanol cleaning and acid-bath cleaning. Ethanol-cleaned microscope slides for PDMS adhesion studies were submerged and spaced apart (not stacked) in a beaker of 200 proof ethanol for at least one day. After soaking, the glass slides were removed and placed on end on absorbent paper to remove excess ethanol and allowed to dry. Acid-cleaned microscope slides were prepared by immersing the glass slides in concentrated sulfuric acid (> 90% w/w) overnight, minimum. For convenience, the slides were kept in a histology rack during acid bath cleaning. Note of caution: Concentrated sulfuric acid is caustic and hygroscopic; leave ample empty vessel volume for fluid expansion. After a minimum of 24 h in sulfuric acid, the histology rack with slides was removed and rinsed with a direct stream of MilliQ DI for 5 min. Clean glass slides were turned on end and dried. For cell culture, acid-cleaned coverslips (Corning, thickness no. 1.5) were prepared following this same cleaning protocol.

### Elastomer Bonding

Immediately after clean glass slides and coverslips were dry, unextracted and E-PDMS plugs, or microchannels, were placed in conformal contact with the glass substrate for thermal bonding using an autoclave. In a comparative study, ultra-violet light (UV) or a dry oven were used for bonding PDMS to glass substrates; high-powered UV with continuous air flow was used to oxidize/activate the PDMS for bonding followed by a 70 °C annealing process. The PDMS glass assemblies were placed in an aluminium foil-lined metal pan, covered and sealed with aluminium foil, and autoclaved with the following settings: 121 oC and 110 kPa for 20 min (sterilization step) with a 20-min drying time (81 °C to 91 °C) (18). After thermal bonding, E-PDMS microfluidic channels for cell culture were cooled, removed from foil in a sterile biosafety hood, transferred to a culture dish and prepared for cell culture.

### Force Measurements

Force measurements of adherent unextracted and E-PDMS structures on glass or PDMS substrates were taken at room temperature after autoclaved samples cool to room temperature (≥ 30 min). Spring force meters (SI Manufacturing) were used for measuring the force required to separate PDMS plugs from substrates. Meter ranges were in Newtons (N) 0-2.5 N, 0-5 N and 0-30 N, the footprint of the PDMS plug is 30.7 mm^2^, results are presented in kPa (1N mm^-2^). Force meters were connected to the PDMS plugs using a strong wire triangle. The triangle base passed through the PDMS plug with the apex attached to the hook of the force meter (Figure 1B). Force meter performance was verified with calibrated weight standards.

### Flow Control and Manipulation

Dynamic fluid manipulations were achieved by modulating positive and negative pressures at the fluidic ports and reservoirs. Culture perfusion was created with differential hydrostatic pressures and gravity flow. For hydrostatic pressures, fluid droplets were placed at the inlet ports or reservoirs to create positive pressures with surface tension. After surface tensions equilibrated, the dish was tilted a few degrees to elevate one end of the coverslip and reinitiate flow. Negative pressures were obtained with unequal concave menisci at the inlets or reservoir. For pulses of stimuli, rapid fluidic infusion was achieved by combining positive pressures (droplets at inlets) with negative pressures (concave menisci at the outlet reservoir), and the infusate was rapidly reversed by wicking the inlet(s) empty with a Kimtech wipe. Re-initiation of flow was achieved by pipetting defined volumes back into the empty inlet. Imaging controls (with flow of fluorescein and fluorescently labelled antibodies) were performed to refine the pulse process. During glial stimulation, the glial culture channel was always maintained at a greater positive pressure than stimulation channels to prevent stimuli from entering the glial culture compartments. Fluidic manipulations were greatly assisted by monitoring experimental duration during image acquisition and utilizing time-mark features in the Zeiss image acquisition software.

### Glial Cell Culture

Animal procedures were conducted in accordance with PHS Policy on Humane Care and Use of Laboratory Animals under approved protocols established through the University of Illinois at Urbana-Champaign Institutional Animal Care and Use Committee. Astrocytes were isolated from P2-P4 Long-Evans BluGill rats through the following procedure: Subjects were rapidly decapitated, the brain removed, meninges cleared from tissues, and the hippocampi and cortex dissected. For each culture, the tissue was minced and incubated with papain (25.5 U/mL, Worthington) or trypsin EDTA (0.05%) for 30 min at 37 °C. After enzymatic digestion, tissue fragments were rinsed with media (without enzyme), and the tissue triturated through a fire-polished glass Pasteur pipette. The cell suspension was centrifuged at 1400 rpm for 5 min and suspended in astrocyte culture media (DMEM with 10% FBS, 3mM L-glutamine, and 100 U/mL penicillin and 0.1mg mL^-1^ streptomycin) and plated at 300-500 cells/mm^2^. Cultures were maintained in astrocyte culture media in a humidified incubator with 5% CO_2_ and 95% air until reaching confluence (~7-9 days). Cultures were shaken at ~300 rpm for four consecutive periods of 18 h to remove unwanted, loosely-adherent cells (i.e., microglia); each 18-h interval was interrupted by ~30-h periods of overnight recovery.^55,56^ Astrocytes were released from the dish and loaded into microfluidic channels where they were allowed to grow and extend branches to compartmentalize into adjacent channels.

### Calcium Imaging

Imaging spatial dynamics of transient Ca^2+^ signals of individual glial cells and networks was achieved using Fluo-4 AM, a cell-permeable Ca^2+^ indicator dye. Cultures were mounted on a Zeiss LSM-510 Meta NLO laser-scanning microscope without an environmental chamber and imaged at room temperature (22-25 °C). Glial cell Ca^2+^ transients were evoked with ATP (20 μM) or glutamate (10 μM). Fluorescence signals were obtained using argon laser illumination (488 nm, 0.1%) through a plan-apochromatic 20x (0.8) objective, and a LP505 filter. Images were acquired with a photomultiplier tube; detector and scanner parameters (gain, sensitivity, scan rate, zoom and field size) were optimized to minimize laser exposure to live cells, and maximize scan rates for field size and resolution. Profiles of Ca^2+^ transients were aligned (from bottom to top of chart) based on the first occurrence Ca^2+^ transients that affected multiple ROI after stimuli markers; transients of individual ROI may be attributed to spontaneous Ca^2+^ spikes.

### Immunocytochemistry

Immunochemistry was performed following our previously published process.^19^ Glial cultures were fixed for 30 min with 4% paraformaldehyde and then permeabilized for 10 min with 0.25% Triton in PBS and blocked for 30 min with 10% BSA in PBS at room temperature. To label glial cells, mouse monoclonal primary antibodies against glial fibrillary acidic protein (GFAP, 1:1000 dilution) were used; antibody-labelling occurred for 1-2 h at room temperature or overnight at 4 °C. Secondary antibodies were goat-anti-mouse Alexa 488 (1:1000 dilution, Molecular Probes) incubated at room temperature for 1–2 h. Rhodamine-conjugated phalloidin (5 U/ml, Molecular Probes) was used to label actin filaments (f-actin) with 20 min incubation at room temperature. The same protocol was followed for immunochemistry in microfluidic channels with flow maintained throughout the labelling process and flow velocities kept to a minimum. Flow directions were reversed every 10 min during the course of the antibody and phalloidin incubations. Following ~20 min PBS rinse, the samples were briefly rinsed with DI water and dried. Channels were filled with Prolong Gold anti-fade reagent (Molecular Probes). To optimize imaging, cells in channels were imaged immediately following antifade reagent application.

## Conclusions

This work advances the fabrication and implementation of microfluidics, particularly for biological applications and cell signalling studies. With conventional equipment (autoclave and solvent hood) available in biological departments, elevated humidity and temperature can be used to induce high-strength bonds between E-PDMS and glass to permit highly biocompatible microfluidics for maintaining fluidic fidelity during growth and stimulation studies. An advantage of this annealing process is that humidified cell culture conditions (37 °C) favour the E-PDMS bonding process, rather than counteract it.

This simple, effective fabrication method will expand the range of possibilities for process miniaturization and sample manipulation in tightly-bound, biocompatible, PDMS-based microfluidics. Solvent extraction or autoclaving alone improves material biocompatibility. Combining these easy processes in sequence retains material biocompatibility while increasing material adhesion, thus improving the versatility of applications of E-PDMS for microfluidic platforms. It is yet to be determined how E-PDMS microfluidics will be advantageous for chemical synthesis, material interactions, flexible electronics, or PDMS surface modifications, but the implications from PDMS deformation from oligomer translocation may prove valuable.

Oligomer translocation between PDMS types is evident through material deformation when oligomer-free, E-PDMS is placed in contact with oligomer-containing PDMS and subjected to elevated temperatures. This provides a possible approach for optimizing curing agent and pre-polymer ratios, and curing conditions to minimize oligomer translocation through, and out of, PDMS.

Optimal cell signalling results when environmental confounds are eliminated. E-PDMS microfluidics are devoid of free oligomers, which are known to accumulate in cells and modify gene transcription. E-PDMS microfluidic platforms are advantageous for a wide range of signalling studies for monotypic cell cultures or co-cultures.

## Acknowledgements

We thank Drs. John Schlup and Chris Culbertson at Kansas State University for their insightful discussions on PDMS-to-glass interactions. We gratefully acknowledge financial support from the National Science Foundation (IGERT 0965918, STC EBICS CBET 0939511, and EAGER DBI 1450962) and the National Institute of Health (MH101655). ORNL is managed by UT Battelle, LLC, for the Department of Energy under contract DE-AC05-00OR22725. We thank Ann Benefiel for her help with the manuscript preparation.

